# Comparative analysis of actin visualization by genetically encoded probes in cultured neurons

**DOI:** 10.1101/2022.08.12.503767

**Authors:** Attila Ignácz, Domonkos Nagy-Herczeg, Angelika Hausser, Katalin Schlett

## Abstract

Actin cytoskeleton predominantly regulates the formation and maintenance of synapses by controlling dendritic spine morphology and motility. To visualize actin dynamics, actin molecules can be labelled by genetically fusing fluorescent proteins to actin monomers or using fluorescently tagged actin-binding proteins or single-chain anti-actin antibodies. However, the effects of these labels on the morphology of neurons have not been quantitatively compared yet. In the present study, we analysed Actin-Chromobody-GFP, LifeAct-GFP and EGFP-actin with respect to their effects on actin-related features in mouse cultured hippocampal neurons.

The actin-binding probes LifeAct and Actin-Chromobody showed similar affinity to F-actin, and along with EGFP-actin, were enriched in dendritic protrusions. In contrast to EGFP-actin, neither of these constructs was able to detect subtle changes in actin remodelling between mature mushroom shaped spine and less developed filopodia. None of the compared probes altered filopodial motility compared to control EGFP expression, however, within 24 hours expression, minor changes in dendritic spine morphology and density were visible. Furthermore, while EGFP-actin and LifeAct-GFP expression did not alter dendritic arborization, AC-GFP expressing neurons displayed a reduced dendritic arborization. We therefore conclude that careful consideration of cellular consequences is required before performing experiments with a particular actin labelling probe in primary neurons.

## Introduction

Actin is a key cytoskeletal element in mammalian cells, involved in many cellular mechanisms (Campellone and Welch, 2010). Neurons, in particular, develop actin-rich growth cones during neurite outgrowth and special postsynaptic structures that are shaped by actin, called dendritic spines. These structures are small protrusions on the dendrites, receiving the majority of glutamatergic synaptic inputs in the brain (Yuste, 2010). Spines usually consist of a bulbous head region where the synaptic receptors and the postsynaptic density accumulates, and a thin neck region that separates the head from the dendritic shaft. Connected to synaptic activity, they can change their morphology, which is a phenomenon called structural plasticity (Arellano *et al*., 2007; Kasai *et al*., 2010; Amtul and Atta-Ur-Rahman, 2015; Bosch and Hayashi, 2015). Spines develop from dendritic filopodia, which are thin protrusions on the shaft without an established synaptic connection (Yuste and Bonhoeffer, 2004; Kayser, Nolt and Dalva, 2008; Ozcan, 2017). Structural remodelling of filopodia and spines underlies changes in synaptic strength on the cellular level. Activity dependent alterations in synaptic strength is, in turn, the cellular basis of learning and memory, placing these protrusions in the focus of neuroscience research (Malenka and Bear, 2004; Nägerl *et al*., 2004; Lisman, Yasuda and Raghavachari, 2012; Nishiyama and Yasuda, 2015).

In the spine heads, actin is present in branched filamentous polymers (F-actin) and globular monomers (G-actin) (Hotulainen and Hoogenraad, 2010; Konietzny, Bär and Mikhaylova, 2017). Although straight filaments are also present in the spine head, they are more common in the neck part, or in immature filopodia (Korobova and Svitkina, 2010). Actin remodelling is tightly regulated in connection with synaptic plasticity and is therefore important in learning and memory as well as in neuronal network functions (Schubert and Dotti, 2007; Honkura *et al*., 2008; Rudy, 2015). This is in accordance with the accumulating evidence that abnormal actin regulation in dendritic spines plays an important role in different neurological diseases and thus research of actin regulation in dendritic spines is crucial in understanding these pathologies (Joensuu, Lanoue and Hotulainen, 2018; Pelucchi, Stringhi and Marcello, 2020).

Fluorescent imaging of actin in living cells is a vital tool to investigate cytoskeletal changes in dendritic spines (Koskinen, Bertling and Hotulainen, 2012). A plethora of fluorescent probes is already available, from cell penetrating small molecules that label filamentous actin in every cell, such as SiR-actin or phalloidin (Lukinavičius *et al*., 2014), to genetically encoded fusion proteins that are expressed in individual cells (Belin, Goins and Mullins, 2014; Melak, Plessner and Grosse, 2017). In case of low density or non-overlapping cells, the former, general actin labelling can be useful, while, among the intermingling network of neurites in neuronal tissue or cell cultures, it is often more desirable to label only a limited number of neurons.

Expression of fluorescently labelled actin monomers is a widely used method in neuronal actin imaging studies (Star, Kwiatkowski and Murthy, 2002; Hotulainen *et al*., 2009; Koskinen, Bertling and Hotulainen, 2012; Melak, Plessner and Grosse, 2017), however, introducing new actin monomers might imbalance the F-G actin treadmilling, leading to alterations in actin-dependent processes (Koskinen and Hotulainen, 2014). Actin-binding protein probes aim to circumvent this disadvantage by recognizing endogenous actin. Most of these molecules consist of sequences from actin-binding proteins like F-tractin (Johnson and Schell, 2009), Utrophin (Patel *et al*., 2017), and LifeAct (Riedl *et al*., 2008). Among these, LifeAct is the smallest, making it an ideal candidate for endogenous actin labelling without interfering with its regulation. An additional but less investigated actin labelling tool is Actin-Chromobody (AC), a small single-chain camelid antibody specific for actin (Rocchetti *et al*., 2014; Traenkle and Rothbauer, 2017).

Both AC and LifeAct, fused to fluorescent proteins, have been successfully used in experiments with living neurons (Belin, Goins and Mullins, 2014; Panza *et al*., 2015; Patel *et al*., 2017; Wegner *et al*., 2017). However, LifeAct-GFP has been reported to have negative effects on various cell types (Flores *et al*., 2019; Xu and Du, 2021), and to exclude certain actin structures from labelling (Munsie *et al*., 2009). This could be explained by the competition of LifeAct with other actin-binding regulatory proteins (Belyy *et al*., 2020). In addition, to date, it has not been thoroughly tested how AC-GFP expression might perturb endogenous actin regulation (Melak, Plessner and Grosse, 2017). Moreover, neither of these probes has been quantitatively compared and analysed in regard to their effects on dendritic branching, spine morphology and motility, despite that these actin-dependent neuronal features predominantly determine neuronal network development and memory functions (Konietzny, Bär and Mikhaylova, 2017).

Therefore, in the present study, we used AC-GFP and LifeAct-GFP as nanobody-based and actin-binding protein-based actin-probe examples, respectively, to test their effects in comparison to the gold standard, EGFP-actin, on actin-related neuronal features. We report that AC-GFP and LifeAct-GFP have similar affinity for actin in spines and show somewhat different relative enrichment in dendritic spines, but only EGFP-actin can reveal small differences in actin dynamics within motile filopodial, and more mature mushroom shaped dendritic protrusions. All actin-labelling probes leave short-term filopodial motility intact. However, overexpression of AC-GFP for 24 hours results in reduced dendritic arborization and increased filopodial density, whereas LifeAct expression induces elongation of dendritic spines. Therefore, we conclude that actin labelling probes must be carefully selected to study neuronal features, because results can vary considerably depending on the functional assay chosen.

## Results and Discussion

### EGFP-actin expression reveals differences in actin remodelling between thin and mushroom spines

To quantitatively compare the three different actin probes, we prepared hippocampal neurons from CD1 wild type murine embryos and let them develop for 12-13 days in vitro (DIV). By this age, cultured neurons extend their neurites, establish synaptic connections and modify their dendritic spines according to cellular and network activity (Dotti, Sullivan and Banker, 1988; Yuste, 2010).

We transfected the cells with plasmids encoding EGFP tagged actin (EGFP-actin), Actin-Chromobody (AC-GFP), or LifeAct (LifeAct-GFP). Besides the actin labelling constructs, mCherry was also co-expressed, providing the possibility to detect spine shape. 24 hours following transfection, time-lapse recordings were taken in every second (Figure 1A, Supplementary Video 1) to reveal fluorescence in dendritic spines. As both AC-GFP and LifeAct-GFP label actin in a non-covalent manner, we first tested their relative affinity to actin, using the FRAP method in cells with a stabilized actin cytoskeleton.

**Figure 1.**
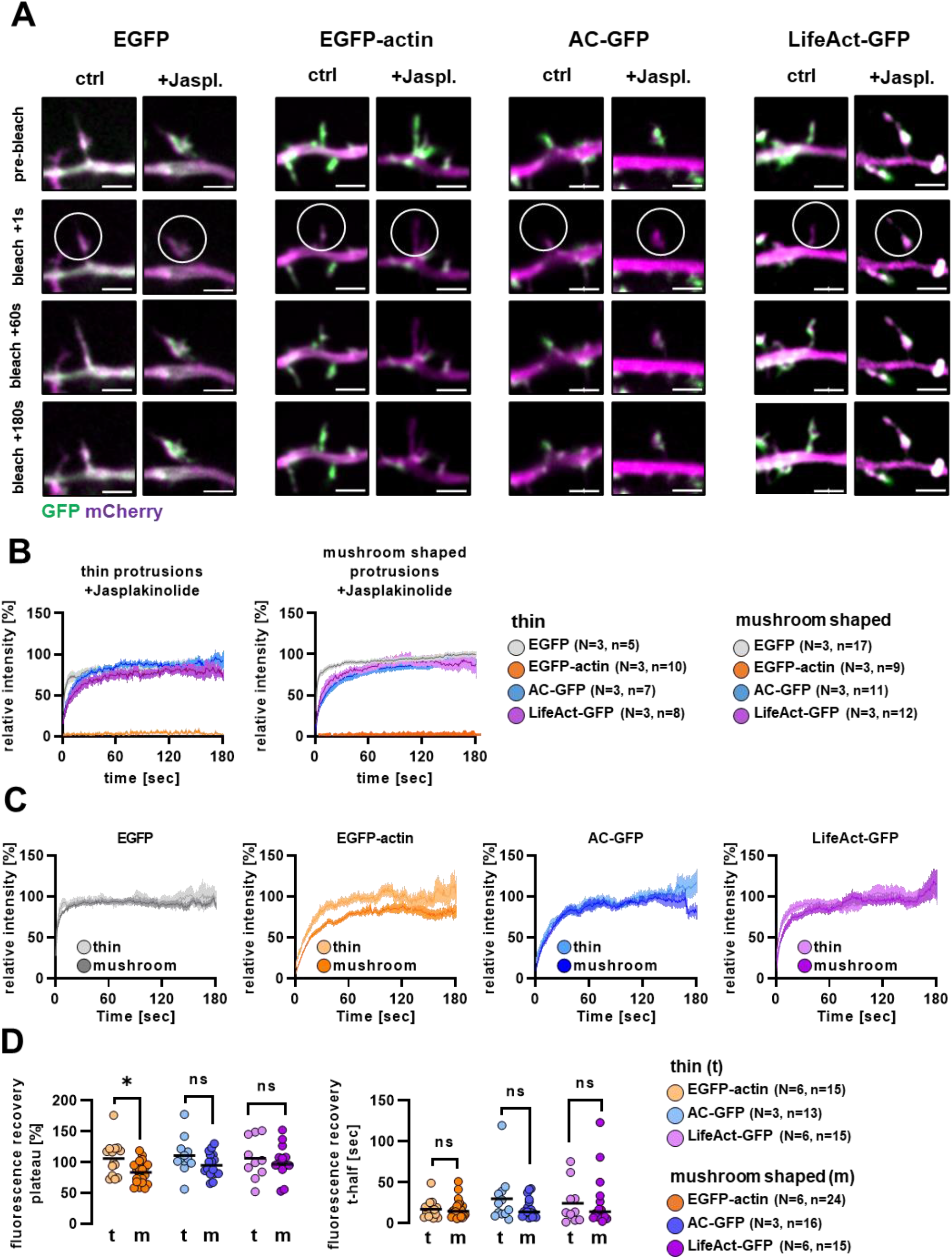
GFP fluorescence recovery after bleaching in cells expressing different actin labelling fusion proteins. (A) Representative live cell confocal image series of cells expressing EGFP, EGFP-actin, Actin-Chromobody-GFP or LifeAct-GFP together with mCherry. 5 µM Jasplakinolide was applied for 7-10 min before imaging. Scalebar: 2 µm. Exported videos are displayed as Supplementary Video 1. Fluorescence recovery curves after photobleaching in cells treated with 5 µM Jasplakinolide (B), and (C) in control cells. Fluorescence was measured in the mCherry positive area, and normalized to the mean of the last 10 pre-bleach values in every spine. Thin and mushroom shaped morphotypes were distinguished based on apparent spine morphology. (D) Fluorescence recovery plateau and t-half values of the analyzed spines. Each point represents an individual spine. N: number of independent cultures, n: number of analysed spines. *p<0.05.

To this end, we left the neuronal cultures untreated or applied Jasplakinolide, an actin stabilizing drug, prior and during imaging, and then bleached the GFP signal within randomly picked dendritic spines (see circled areas in Figure 1A). Although Koskinen and colleagues reported that mushroom shaped spine heads increase in size 300 seconds after photobleaching of EGFP-actin (Koskinen and Hotulainen, 2014), we did not find differences in spine diameter between pre-bleach and 180sec post-bleach values in either control or Jasplakinolide treated groups (Supplementary Figure 1A).

Both thin and mushroom shaped protrusions were bleached, then the two groups were analysed separately (Figure 1B and 1C). EGFP expressing cells served as negative control as the fluorescent protein itself lacks actin-binding affinity. EGFP-actin, on the other hand, served as positive control, as it incorporates into the endogenous F-actin. In Jasplakinolide-treated neurons, as expected, we observed an almost immediate recovery of fluorescence in the EGFP expressing group, whereas the EGFP-actin signal did not recover. AC-GFP and LifeAct-GFP signals, on the other hand, reappeared within the bleached protrusions (Figure 1A, Figure 1B, Supplementary Video 1).

To see how reliably the actin probes studied can track subtle changes in actin dynamics we analyzed FRAP in untreated control neurons. As before, EGFP recovered almost immediately, preventing recovery curve analysis. In case of the actin probes, one-phase decay curves were fitted to individual fluorescence recovery data, then recovery half-time (t-half) and plateau values were extracted, representing the speed of recovery and the mobile actin fraction, respectively (Figure 1C). Because mushroom-shaped spines have a stable actin cytoskeletal core within the head region (Hlushchenko, Koskinen and Hotulainen, 2016), we assumed that thin and mushroom shaped protrusions differ in their ratio of mobile actin. Indeed, in EGFP-actin expressing cells, plateau values were significantly smaller in mushroom than in thin spines, while recovery half-time values did not differ significantly. In contrast, neurons expressing AC-GFP or LifeAct-GFP had indistinguishable recovery curves between mushroom-shaped and thin protrusions (Figure 1C, Figure 1D).

Our results are in accordance with FRAP data obtained with F-tractin (Johnson and Schell, 2009), showing that fluorescence recovery of both actin-binding protein-based and nanobody-based probes is independent from actin remodelling and depends only on their affinity to F-actin, that is similar in AC-GFP and LifeAct-GFP (Figure 1A, Figure 1B, Supplementary Video 1). In addition, our results suggest that even without photomanipulation, minor changes in actin reorganization cannot be revealed when using actin-binding probes.

### Genetically encoded actin labelling probes do not alter filopodial motility

Dendritic filopodia are small filamentous structures whose protrusion and retraction are driven predominantly by their actin cytoskeleton (Kayser, Nolt and Dalva, 2008; Hotulainen and Hoogenraad, 2010; Ozcan, 2017). Therefore, we investigated how the different actin probes affect the dynamics of filopodial motility, by comparing the average displacement of intensity weighted center of mass of selected filopodia (Figure 2A, Supplementary Video 2).

**Figure 2.**
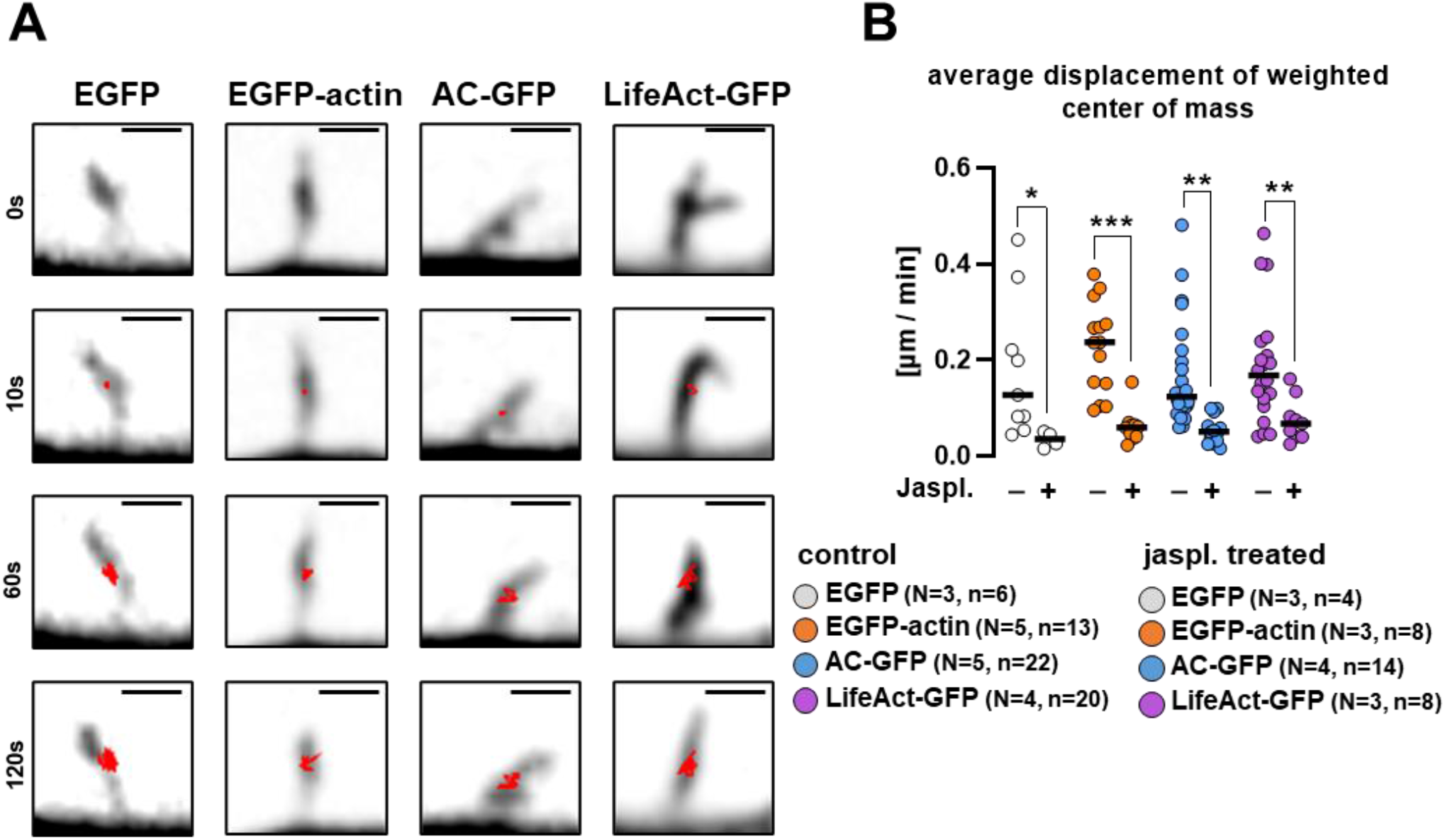
Dendritic filopodial motility in cells expressing EGFP, EGFP-actin, Actin-Chromobody-GFP, or LifeAct-GFP. (A) Representative inverted mCherry fluorescence confocal image series of filopodia. Red lines show the movement track of intensity weighted center of mass. Scalebar: 1 µm. Exported video is presented in Supplementary Video 2. (B) Average intensity-weighted center of mass displacement over 60s time periods in control vs 5 µM Jasplakinolide treated cells. Each point represents an individual spine. Black lines show medians. Numbers of elements: N: independent cultures, n: analysed filopodia *p<0.05 **p<0.01 ***p<0.001.

We found no significant difference in center of mass displacement between actin probe expressing cells and EGFP expressing control filopodia (Figure 2B), even when the filopodia were bleached (see previous FRAP experiments and Supplementary Figure 1B). As expected, Jasplakinolide treatment completely blocked motility in every transfected protrusion (Figure 2B). Our data suggest that the tested actin labelling proteins do not interfere with fast morphological changes in these temporal protrusions.

### Actin-labelling probes exert distinct effects on dendritic spine morphology and density

FRAP experiments already indicated an enrichment of actin labelling probes in dendritic protrusions, in accordance with the elevated amount of F-actin in spines (Korobova and Svitkina, 2010). Thus, we systematically quantified the relative spine-to-shaft GFP fluorescence distribution in live cells (Figure 3A). EGFP spine-to-shaft fluorescence ratio served as a control, as it is determined by the difference in thickness between shaft and spine. As expected, we observed a significant increase in spine-to-shaft ratio in all the actin probe expressing cells compared to EGFP expression, suggesting that actin probes were enriched in F-actin rich dendritic spines. In the case of AC-GFP, our results are in accordance with a study by Wegner and colleagues (Wegner *et al*., 2017), although in non-neuronal cell types, this probe was reported to show a high fluorescence background (Melak, Plessner and Grosse, 2017). In addition, LifeAct-GFP expressing cells had a significantly lower level of relative enrichment in spines compared to EGFP-actin and AC-GFP (Figure 3A, Figure 3B – see the elevated LifeAct-GFP fluorescence intensity within the dendritic shaft). This is like due to the known high background fluorescence level of LifeAct, originating from its affinity to G-actin (Melak, Plessner and Grosse, 2017).

**Figure 3.**
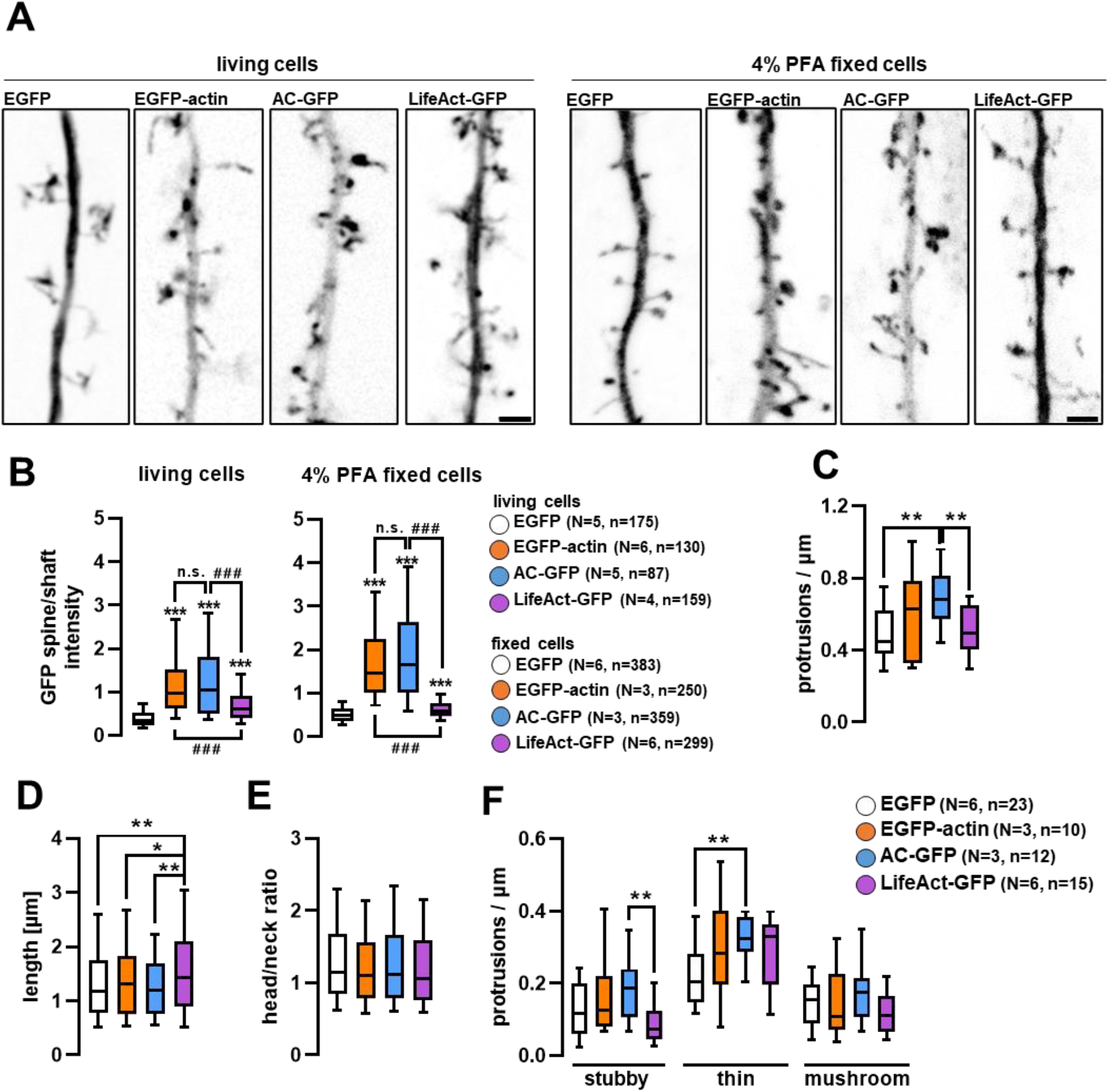
Dendritic spine localization of actin probes and their effect on dendritic spine morphology. (A) Representative inverted GFP fluorescence confocal images of dendritic segments. Scalebars: 2 µm. (B) Relative fluorescence intensity in dendritic spines compared to an adjacent shaft segment in live and fixed cell images. (C) Protrusion density of dendritic segments expressing the actin labelling fusion proteins. (D) Length and (E) head-to-neck ratio of individual dendritic spines of transfected cells. (F) Protrusion density of different morphotypic spines on transfected dendritic segments. Boxes show median and interquartile range, whiskers show 10-90%. Numbers of elements in (C) (D) (E): N: independent cultures, n: analysed dendrites. In (B), N: independent cultures, n: analysed spines. *p<0.05 **p<0.01 ***p<0.001. ### p<0.001. In (B) and (C) *** is compared to EGFP control, ### is compared to the connected groups.

For spine morphometric analysis, fixed cultures were used (Figure 3). Importantly, elevated spine-to-shaft fluorescence ratio of actin probe was further increased after fixation, especially in case of EGFP-actin and AC-GFP (Figure 3B).

Dendritic spine morphology and density were analysed on high resolution confocal images along secondary dendritic branches. Expression of EGFP-actin or LifeAct-GFP for 24 hours did not influence total protrusion density, nevertheless it was significantly higher in AC-GFP expressing neurons (Figure 3C). In addition, we further analysed spine morphology by measuring spine length, neck and head diameters on max-projected images. While EGFP-actin or AC-GFP expression did not alter spine length, LifeAct-GFP expressing spines were significantly longer compared to all other conditions (Figure 3D). At the same time, head/neck ratio was not affected (Figure 3E). Based on these morphometric values, three main morphotypes were distinguished, in accordance with the literature: stubby, thin, and mushroom protrusions (Peters and Kaiserman-abflamof, 1970; Bourne and Harris, 2008; Rochefort and Konnerth, 2012). When the density of the distinct morphotypes was compared, thin protrusion density in AC-GFP cells was significantly elevated in relation to EGFP expressing cells (Figure 3F).

Neither mushroom nor stubby morphotype densities were affected compared to control EGFP expressing neurons. Therefore, the initially observed difference in protrusion density is likely due to the higher density of thin protrusions in AC-GFP expressing cultured neurons, as shown in Figure 3F. Although previous in vivo observations using STED imaging did not reveal striking differences in spine morphology of LifeAct or AC expressing neurons (Wegner *et al*., 2017), our study provides quantitative data showing that overexpression of AC-GFP and LifeAct-GFP can indeed lead to subtle changes in spine morphology and density.

### Actin-Chromobody expression impairs dendritic arborization within 24 hours

Besides dendritic spine morphology, we also investigated the intracellular distribution of the tested actin-probes within transfected neurons. As expected, all constructs were present in both the somatodendritic and axonal compartments (Figure 4A – lower cut-outs highlight the indicated axonal segments). We did not observe striking differences in axonal or growth cone morphology between the different groups (data not shown).

**Figure 4.**
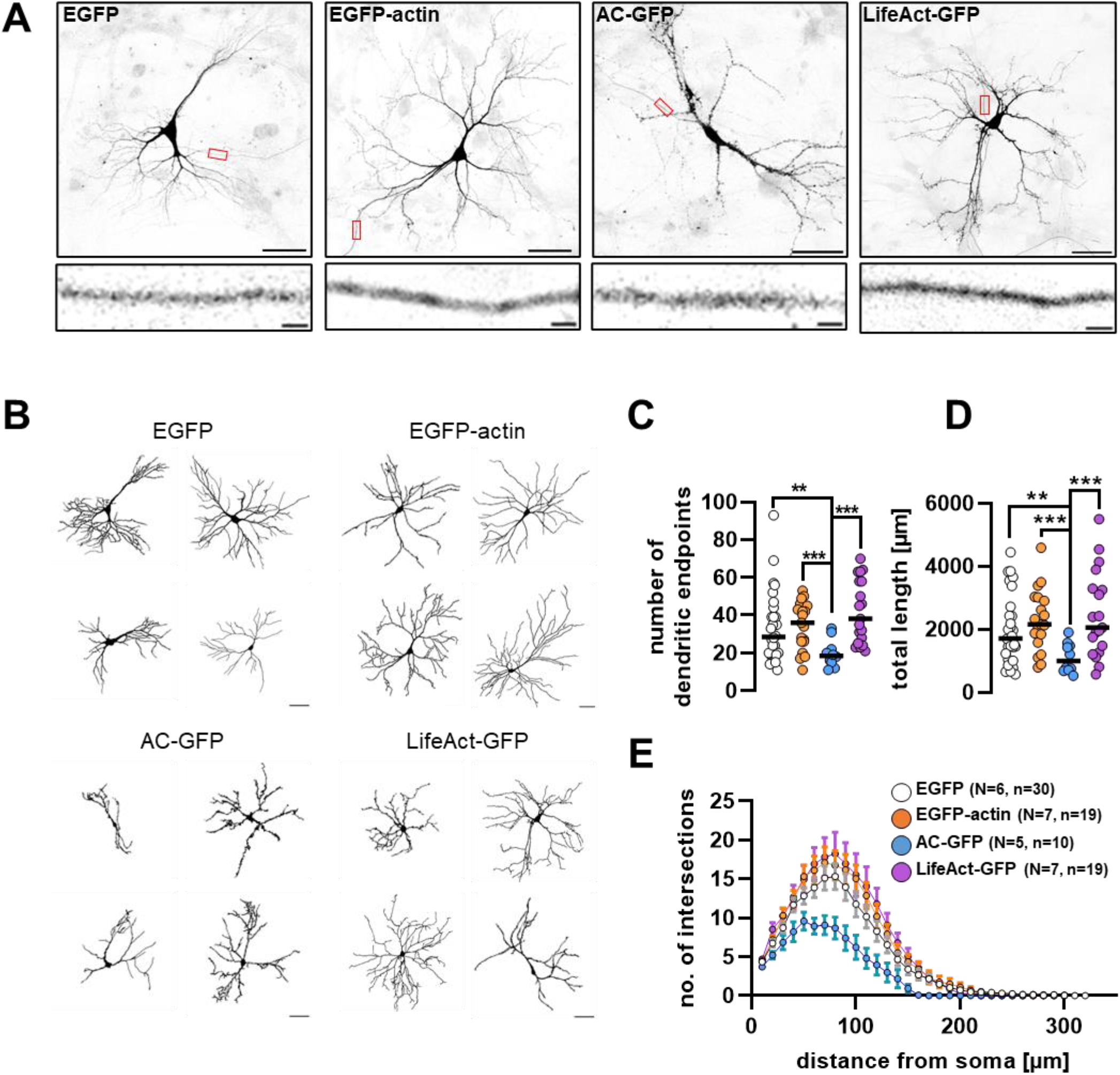
Cellular distribution of actin labelling fusion proteins and their effect on dendritic morphology. (A) Confocal images of cultured hippocampal neurons expressing actin labelling fusion proteins. The images are shown in inverted GFP fluorescence. Scalebar: 50 µm. Cut-out axonal parts are indicated by red rectangles. Scalebar: 2 µm. (B) Representative dendritic outlines used for morphometric analysis. Scalebar: 50 µm. (C) Number of dendritic endpoints and (D) total dendritic length of neurons expressing the different fusion proteins. Each data point presents an individual cell, black lines show median. (E) Number of dendritic intersections with increasing radius circles starting from the soma. Data presented as mean +-s.e.m. Numbers of elements: N: independent cultures, n: individual cells. **p<0.01 ***p<0.001.

As Patel and colleagues reported that long-term expression of certain actin-binding protein-based probes in developing neurons influence dendritic morphology (Patel *et al*., 2017), we analysed whether dendritic arborization is altered within 24 h after the transfection of the tested probes. Confocal images were quantitatively evaluated using Ilastik machine learning program to segment dendritic trees and dendritic endpoints (Berg *et al*., 2019). Examples of the analysed dendritic outlines from each tested group are shown in Figure 4B. According to our data, neither EGFP-actin nor LifeAct-GFP expression altered dendritic endpoint number, total dendrite length or the extent of dendritic arborization when compared to the EGFP control (Figure 4C-E), consistent with the existing literature (Belin, Goins and Mullins, 2014; Patel *et al*., 2017).

On the other hand, AC-GFP expressing cells had significantly fewer dendritic endpoints (Figure 4C) and a shorter total dendritic length (Figure 4D) than control, EGFP expressing neurons. In addition, AC-GFP expressing neurons had significantly shorter dendrites and fewer branches compared to the other constructs (Figure 1E). This may seem to contradict another study, which reported that zebrafish embryos can express this fusion protein at various stages of development without apparent impairment of cell function (Panza *et al*., 2015; Melak, Plessner and Grosse, 2017). However, in the absence of such detailed analysis of neuronal morphology, altered dendritic arborization in fish embryos cannot be excluded. Furthermore, to our knowledge, our report is the first to quantitatively compare dendritic morphology of AC expressing neurons to control ones, and in the present study we provide evidence that AC expression can reduce dendritic arborization already within 24 hours.

Overall, our results support the view that expression of genetically encoded actin probes is generally an appropriate mean to visualize actin cytoskeleton structures in individual neurons. However, we found specific limitations in using both nanobody (AC) and actin-binding protein-based (LifeAct-GFP) probes in cultured neurons (Table 1), highlighting the importance of careful testing the cellular consequences before conducting experiments with any given actin labelling probe.

**Table 1.**
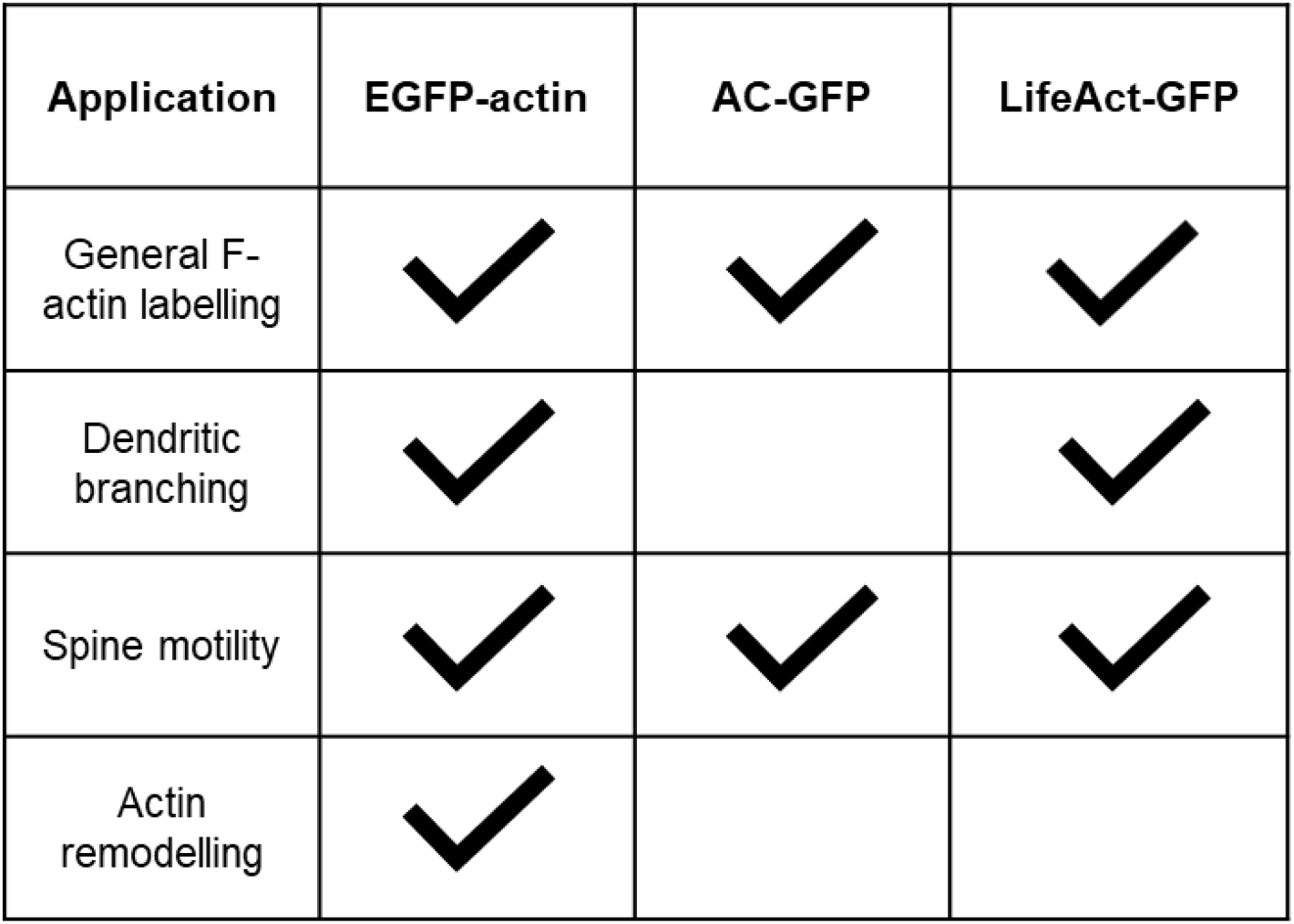
Suitability of the compared actin labelling fusion proteins in different fields of application. Check mark means endorsement based on our results.

## Materials and Methods

### Animal handling

Wild type CD1 mice were housed in the animal facility at 22 ± 1°C with 12-h light/dark cycles and ad libitum access to food and water. All experiments complied with local guidelines and regulations for the use of experimental animals, in agreement with European Union and Hungarian legislation.

### Cell cultures

Embryonic hippocampal cultures were prepared from CD1 mice on embryonic day 17-18, according to (Morales *et al*., 2021). Cells were seeded onto poly-L-lysine (Sigma-Aldrich #P5899) and Laminin (R&D Systems #3446-005-01, 4-8 µg/cm^2^) coated glass-bottom Petri dishes (Greiner Bio-One #627975) or onto PLL-Laminin coated glass coverslips (Marienfeld-Superior #0111520) in 24 well plates (Greiner Bio-One #66210). Glass coverslips had been previously plasma cleaned for 5 min each side using O_2_ plasma. Cell density was 1.4 × 10^5^ cells per coverslip or compartment. The cultures were maintained for 12-13 days in vitro in 5% CO_2_ at 37°C before experiments.

The cells were initially seeded in NeuroBasal PLUS (ThermoFisher Scientific #A35829-01) culture medium supplemented by 2% B27 PLUS (ThermoFisher Scientific #A3582801), 5% foetal bovine serum (PAN Biotech #P30-3309), 0.5 mM GlutaMAX (ThermoFisher Scientific #35050-038) 40 μg/mL gentamycin (Sigma, #G1397) and 2.5 μg/mL Amphotericin B (Gibco #15290-026). On the 5^th^ day *in vitro* (DIV), half of the medium was changed to BrainPhys (StemCell Technologies #05790) supplemented with 2% SM1 (StemCell Technologies #05711), 40 μg/mL gentamycin and 2.5 μg/mL Amphotericin B medium. On DIV9, one-third of the medium was again changed to BrainPhys supplemented with SM1, 40 μg/mL gentamycin and 2.5 μg/mL Amphotericin B medium. Depending on glial confluency, on DIV4-6, 10µM CAR (cytosine-arabinofuranozide (Sigma-Aldrich #C6645)) was added to the cultures.

### Transfection and chemical treatment

Transfection was carried out on DIV12-13 using Lipofectamine STEM reagent (ThermoFisher Scientific STEM00001) according to the manufacturer’s guide. The following plasmid constructs were used: pmCherry-N1 (Clontech), pEGFP-N1 (Clontech), pEGFP-actin (Clontech), Actin-Chromobody-TagGFP2 (AC-GFP, Chromotek), LifeAct-TagGFP2 (Clontech). Each GFP-conjugated protein was co-expressed with mCherry to highlight neuronal morphology. 24 hours after transfection, cells were either fixed or used for live cell imaging.

Jasplakinolide (HelloBio #HB3946) was used in a final concentration of 5 μM on glass-bottom Petri dishes mounted for live cell imaging, 7-15 min before imaging, and was present throughout the imaging session.

### Confocal microscopy in fixed samples

The cultures were fixed using 4% PFA in PBS for 20 min, followed by 3 × 5 min PBS wash. Then, the coverslips were mounted in ProLong Diamond Antifade mountant (Invitrogen #P36961).

Images were taken with a Zeiss LSM 800 microscope with a Plan-Apochromat 20x/0.8 dry objective for the dendritic tree analysis, and with a Plan-Apochromat 63x/1.4 Oil objective for the dendritic spine density analysis. Z-stack images were taken in 0.7 μm and 0.2 μm intervals, respectively.

### Live cell imaging and FRAP experiments

For FRAP experiments, the culture medium was changed to a custom-made imaging buffer containing 142 mM NaCl, 5.4 mM KCl, 1.8 mM CaCl_2_, 1mM NaH_2_PO_4_, 0.8mM MgSO_4_, 5mM glucose and 25 mM HEPES with a pH of 7.4. Time-lapse images were taken with a Zeiss Axio Observer microscope equipped with a CSU-X1 Spinning Disk module, using a Plan-Apochromat 100x/1.46 Oil objective. Selective photobleaching of the GFP signal was done by a separately controlled RAPP UGA42 laser, emitting 473 nm laser beam. Images were taken every second for 360 seconds. After the first 10-20 images were taken, 5-10 circular spots were bleached separately. Fluorescence recovery was recorded for at least 180 sec post-bleach in every case (Supplementary Video 2).

### Image analysis methods

Dendritic branching was analysed using FIJI and Ilastik softwares. Z-stack images were max-projected and a two-stage Ilastik random forest model was trained to segment the foreground (the neuronal dendritic tree) and background in the pictures, followed by manual correction. A second Ilastik model was trained to segment the binary images further, adding dendritic skeleton and endpoints. Sholl analysis was carried out on the segmented images, using the built-in FIJI function. Dendritic endpoints were counted, and image skeletons were analysed also by the corresponding FIJI functions (Supplementary Figure 2A).

Dendritic spine densities and morphotypes were analysed on max-projected z-stack images as well. Here, Ilastik models were trained to segment first the dendrites and background, second the intra-spine components: base, neck, head. A separate third model was trained to draw a line inside the spines and the dendrite so their running length could be measured. Spine density and morphotype data were then summarized in FIJI: dendritic length was measured, spines counted, their length, base dimeter and head diameter measured (Supplementary Figure 2B). Three morphotypes were distinguished: stubby spines that were shorter than 0.8 μm, thin spines that were longer than 0.8 μm and their head/neck diameter ratios were smaller than 1.5, and mushroom spines that were longer than 0.8 μm and head/neck ratio higher than 1.5. The ratios used for morphotyping were according to (Morales *et al*., 2021).

Spine-to-shaft ratio of GFP fluorescence was measured on max-projected z-stack images that had been used for spine density measurements in fixed cells, or the first frame of live cell recordings. In both cases Ilastik models were trained to segment background, shaft, and spine. The spines were then enlarged as circles and taken as regions of interest (ROI). Spine pixels in the ROI were used as mask for spine intensity, shaft pixels for shaft intensity, and background pixels for background intensity (Supplementary Figure 2C). Hence, the spine intensities were normalized to and adjacent dendritic segment only instead of the full branch which had visible heterogeny in intensity, as well as in the case of background which was also heterogenous.

2-channel FRAP recordings were analysed with a custom-made FIJI plugin called FRAP2ch the following way: in the bleached spines, mCherry signal was used as a mask to select the region of interest in the GFP channel, which was then measured, in every timeframe. Every spine intensity was normalised to the intensity of a nearby dendritic segment, which were both background-corrected. Then, spine intensities were normalized to the mean of the last 10 pre-bleach time points. A one-phase decay equation was then fitted to the post-bleach values, according to (Koskinen and Hotulainen, 2014). T-half and plateau values were calculated for each spine individually, using GraphPad Prism.

To measure dendritic spine motility, another custom-made FIJI plugin, called DFMA was used (Tárnok *et al*., 2015). Here, the mCherry signal was used on filopodial protrusions to locate their intensity-weighted center of mass on every timepoint. Then, cumulative displacement curves were calculated, and the 60 sec points were compared and statistically analysed (Supplementary Video 1).

## Statistical analysis

Statistical analysis of the acquired data was carried out using Graphpad Prism software 6. First, Shapiro-Wilk test was used to determine normality of distribution for each dataset. Then, comparison was done depending on the normality: for datasets showing Gaussian distribution One-Way ANOVA tests were carried out and Kruskal-Wallis test for those showing nonparametric distribution.

## Supporting information

Ignacz et al_Supplementary video_1

Ignacz et al_Supplementary video_2

## Acknowledgements

This work was supported by a grant from the German Research Foundation (DFG HA-357/11-3) to A.H. and a travel exchange program by the DAAD (PPP Hungary 57392635 and 57602951; 449419 and 169036 by TKA) to A.H. and K.S., respectively, as well as by the Hungarian Brain Research Program (2017-1.2.1-NKP-2017-00002) and by the VEKOP-2.3.3-15-2016-00007 grants. A.I. and D.N-H. are grateful for the support of the ÚNKP-21-3 New National Excellence Program of the Ministry for Innovation and Technology from the source of the National Research, Development and Innovation Fund, and National Talent Program NTP-NFTÖ-21-B-0250, respectively.

---

**Supplementary Figure 1.**
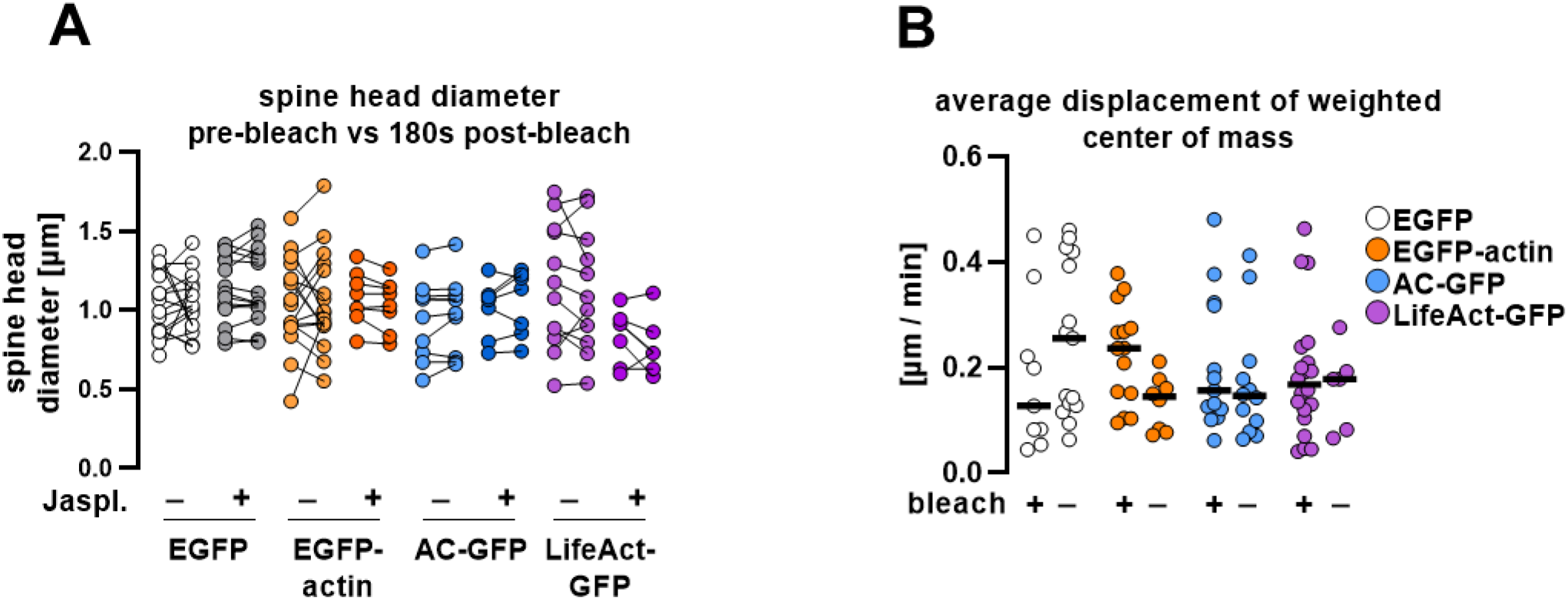
(A) Spine head diameter changes of transfected and GFP-photobleached cells, in regard to 5 µM Jasplakinolide treatment. Data are represented as before-after, where before is the pre-bleach value, after is bleach + 180 sec value. Each point shows one spine. (B) Filopodial motility in unbleached vs photobleached cells. Each point represents an individual spine. Black lines show medians. No significant differences were found.

**Supplementary Figure 2.**
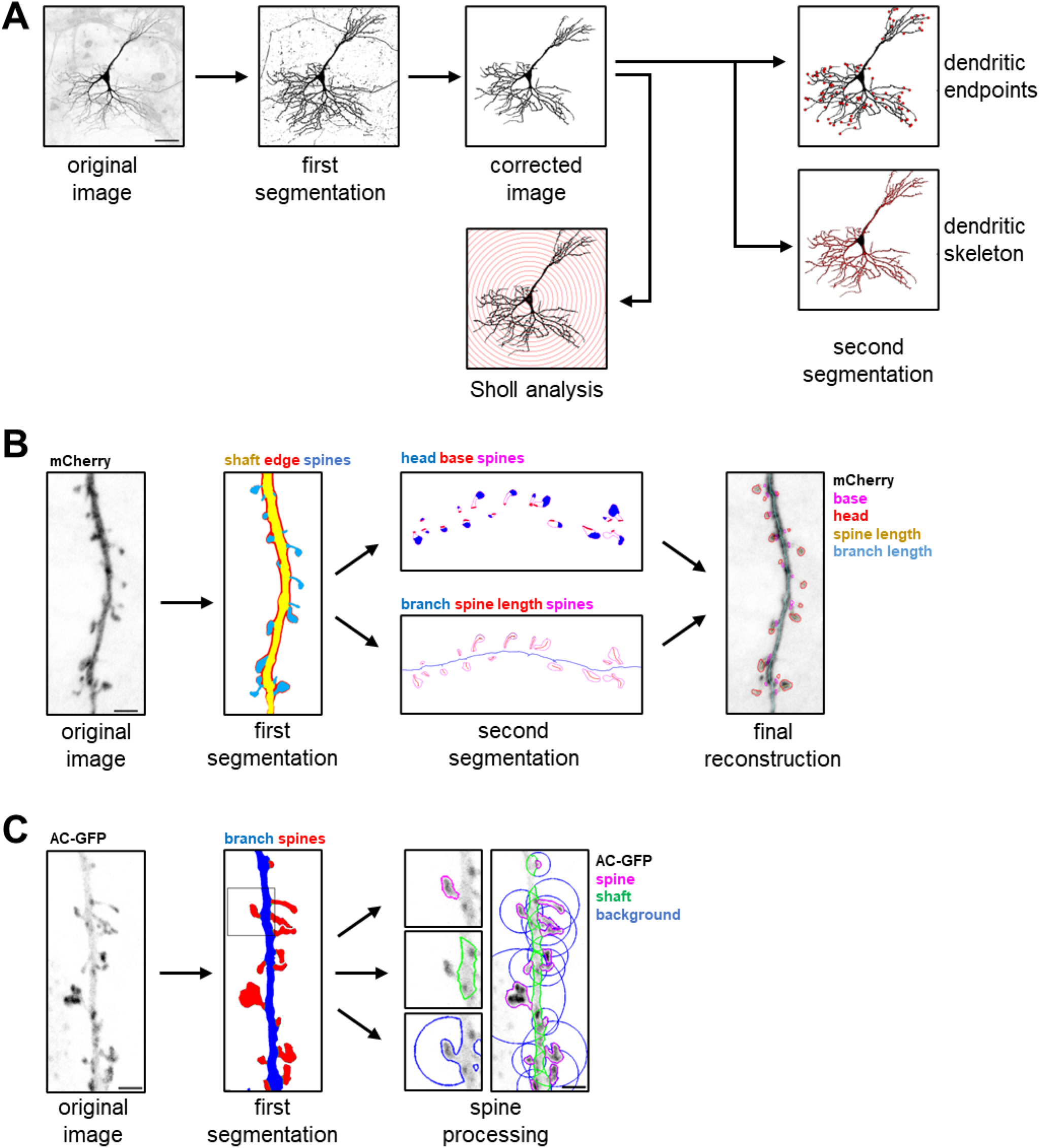
Workflow of machine learning based image analysis methods used in the present work. (A) Sholl analysis, dendritic endpoints and skeleton finding. Original and thresholded images are shown in inverted fluorescence, measured properties are shown in red. Scalebar: 50 µm.(B)Dendritic spine morphometric measurements. Original image is shown in inverted fluorescence, the segmentation categories are written in their corresponding colors in the images. Scalebar: 2 µm.(C)Quantification of fluorescence enrichment in dendritic spines. Original image is shown in inverted fluorescence, the segmentation categories are written in their corresponding colors in the images. Scalebar: 2 µm.

**Supplementary Video 1**. GFP fluorescence recovery after bleaching in cells expressing different actin labelling fusion proteins. Representative live cell confocal time-lapse recordings of GFP (green) together with mCherry (magenta). 5 µM Jasplakinolide was applied for 7-10 min before imaging. Scalebar: 2 µm.

**Supplementary Video 2**. Filopodial motility in cells expressing different genetically encoded actin probes. Representative mCherry fluorescence confocal time-lapse recordings of filopodia. 5 µM Jasplakinolide was applied for 7-10 min before imaging. Red circles show current center of mass in every frame, yellow lines show the movement track. Scalebar: 1 µm.

## References

Amtul, Z. and Atta-Ur-Rahman (2015) ‘Neural plasticity and memory: Molecular mechanism’, Reviews in the Neurosciences, 26(3), pp. 253–268. doi: 10.1515/revneuro-2014-0075.

Arellano, J. I. et al. (2007) ‘Ultrastructure of dendritic spines : correlation between synaptic and spine morphologies’, Frontiers in Neuroscience, 1(1), pp. 131–143. doi: 10.3389/neuro.01/1.1.010.2007.

Belin, B. J., Goins, L. M. and Mullins, R. D. (2014) ‘Comparative analysis of tools for live cell imaging of actin network architecture’, BioArchitecture. Taylor and Francis Inc., 4(6), pp. 189–202. doi: 10.1080/19490992.2014.1047714/SUPPL_FILE/KBIA_A_1047714_SM6287.ZIP.

Belyy, A. et al. (2020) ‘Structure of the Lifeact–F-actin complex’, PloS Biology, 18(11), pp. 1–18. doi: 10.1371/journal.pbio.3000925.

Berg, S. et al. (2019) ‘Ilastik : Interactive Machine Learning for Image Analysis’, Nature Methods, 16, pp. 1226–1232. Available at: https://doi.org/10.1038/s41592-019-0582-9.

Bosch, M. and Hayashi, Y. (2015) ‘Structural plasticity of dendritic spines’, Current Opinion in Neurobiology. Elsevier Ltd, 22(3), pp. 383–388. doi: 10.1016/j.conb.2011.09.002.Structural.

Bourne, J. N. and Harris, K. M. (2008) ‘Balancing Structure and Funcion at Hippocampal Dendritic Spines’, Annu Rev Neurosci, 31, pp. 47–67. doi: 10.1146/annurev.neuro.31.060407.125646.Balancing.

Campellone, K. G. and Welch, M. D. (2010) ‘A nucleator arms race: cellular control of actin assembly.’, Nature reviews. Molecular cell biology, 11(4), pp. 237–51. doi: 10.1038/nrm2867.

Dotti, C. G., Sullivan, C. A. and Banker, G. A. (1988) The Establishment of Polarity by Hippocampal Neurons in Culture, The Journal of Neuroscience.

Flores, L. R. et al. (2019) ‘Lifeact-TagGFP2 alters F-actin organization, cellular morphology and biophysical behaviour’, Scientific Reports 2019 9:1. Nature Publishing Group, 9(1), pp. 1–13. doi: 10.1038/s41598-019-40092-w.

Hlushchenko, I., Koskinen, M. and Hotulainen, P. (2016) ‘Dendritic spine actin dynamics in neuronal maturation and synaptic plasticity’, Cytoskeleton, 73(9), pp. 435–441. doi: 10.1002/cm.21280.

Honkura, N. et al. (2008) ‘The Subspine Organization of Actin Fibers Regulates the Structure and Plasticity of Dendritic Spines’, Neuron, 57(5), pp. 719–729. doi: 10.1016/j.neuron.2008.01.013.

Hotulainen, P. et al. (2009) ‘Defning mechanisms of actin polymerization and depolymerization during Dendritic spine morphogenesis’, Journal of Cell Biology, 185(2), pp. 323–339. doi: 10.1083/jcb.200809046.

Hotulainen, P. and Hoogenraad, C. C. (2010) ‘Actin in dendritic spines: Connecting dynamics to function’, Journal of Cell Biology, 189(4), pp. 619–629. doi: 10.1083/jcb.201003008.

Joensuu, M., Lanoue, V. and Hotulainen, P. (2018) ‘Dendritic spine actin cytoskeleton in autism spectrum disorder’, Progress in Neuro-Psychopharmacology and Biological Psychiatry. Elsevier, 84(September 2017), pp. 362–381. doi: 10.1016/j.pnpbp.2017.08.023.

Johnson, H. W. and Schell, M. J. (2009) ‘Neuronal IP3 3-kinase is an F-actin-bundling protein: Role in dendritic targeting and regulation of spine morphology’, Molecular Biology of the Cell. The American Society for Cell Biology, 20(24), pp. 5166–5180. doi: 10.1091/MBC.E09-01-0083/ASSET/IMAGES/LARGE/ZMK0240992920010.JPEG.

Kasai, H. et al. (2010) ‘Structural dynamics of dendritic spines in memory and cognition’, Trends in Neurosciences. Elsevier Ltd, 33(3), pp. 121–129. doi: 10.1016/j.tins.2010.01.001.

Kayser, M. S., Nolt, M. J. and Dalva, M. B. (2008) ‘EphB receptors couple dendritic filopodia motility to synapse formation’, Neuron. Neuron, 59(1), pp. 56–69. doi: 10.1016/J.NEURON.2008.05.007.

Konietzny, A., Bär, J. and Mikhaylova, M. (2017) ‘Dendritic actin cytoskeleton: Structure, functions, and regulations’, Frontiers in Cellular Neuroscience, 11(May), pp. 1–10. doi: 10.3389/fncel.2017.00147.

Korobova, F. and Svitkina, T. M. (2010) ‘Molecular architecture of synaptic actin cytoskeleton in hippocampal neurons reveals a mechanism of dendritic spine morphogenesis’, Molecular biology of the cell. Mol Biol Cell, 21(1), pp. 165–176. doi: 10.1091/MBC.E09-07-0596.

Koskinen, M., Bertling, E. and Hotulainen, P. (2012) Methods to measure actin treadmilling rate in dendritic spines. 1st edn, Methods in Enzymology. 1st edn. Elsevier Inc. doi: 10.1016/B978-0-12-388448-0.00011-5.

Koskinen, M. and Hotulainen, P. (2014) ‘Measuring F-actin properties in dendritic spines’, Frontiers in Neuroanatomy, 8(AUG), pp. 1–14. doi: 10.3389/fnana.2014.00074.

Lisman, J., Yasuda, R. and Raghavachari, S. (2012) ‘Mechanisms of CaMKII action in long-term potentiation.’, Nature reviews. Neuroscience. Nature Publishing Group, 13(3), pp. 169–82. doi: 10.1038/nrn3192.

Lukinavičius, G. et al. (2014) ‘Fluorogenic probes for live-cell imaging of the cytoskeleton’, Nature Methods, 11(7), pp. 731–733. doi: 10.1038/nmeth.2972.

Malenka, R. C. and Bear, M. F. (2004) ‘LTP and LTD: An embarrassment of riches’, Neuron, 44(1), pp. 5–21. doi: 10.1016/j.neuron.2004.09.012.

Melak, M., Plessner, M. and Grosse, R. (2017) ‘Actin visualization at a glance’, Journal of Cell Science, 130(3), pp. 525–530. doi: 10.1242/jcs.189068.

Morales, C. O. et al. (2021) ‘Protein kinase D promotes activity-dependent AMPA receptor endocytosis in hippocampal neurons’, Traffic, 22(12), pp. 454–470. doi: 10.1111/tra.12819.

Munsie, L. N. et al. (2009) ‘Lifeact cannot visualize some forms of stress-induced twisted f-actin’, Nature Methods, 6(5), p. 317. doi: 10.1038/nmeth0509-317.

Nägerl, U. V. et al. (2004) ‘Bidirectional activity-dependent morphological plasticity in hippocampal neurons’, Neuron, 44(5), pp. 759–767. doi: 10.1016/j.neuron.2004.11.016.

Nishiyama, J. and Yasuda, R. (2015) ‘Biochemical Computation for Spine Structural Plasticity’, Neuron. Elsevier Inc., 87(1), pp. 63–75. doi: 10.1016/j.neuron.2015.05.043.

Ozcan, A. S. (2017) ‘Filopodia: A rapid structural plasticity substrate for fast learning’, Frontiers in Synaptic Neuroscience, 9(JUN), pp. 1–9. doi: 10.3389/fnsyn.2017.00012.

Panza, P. et al. (2015) ‘Live imaging of endogenous protein dynamics in zebrafish using chromobodies’, Development (Cambridge). Company of Biologists Ltd, 142(10), pp. 1879–1884. doi: 10.1242/DEV.118943/VIDEO-6.

Patel, S. et al. (2017) ‘Functional characterisation of filamentous actin probe expression in neuronal cells’, PLoS ONE. Public Library of Science, 12(11), pp. 1–18. doi: 10.1371/journal.pone.0187979.

Pelucchi, S., Stringhi, R. and Marcello, E. (2020) ‘Dendritic spines in alzheimer’s disease: How the actin cytoskeleton contributes to synaptic failure’, International Journal of Molecular Sciences, 21(3), pp. 1–23. doi: 10.3390/ijms21030908.

Peters, A. and Kaiserman-abflamof, I. T. A. R. (1970) ‘The Small Pyramidal Neuron o f the Rat Cerebral Cortex. The Perikaryon, Dendrites and Spines’, Am. J. Anat., 127, pp. 321–355. Available at: https://doi.org/10.1002/aja.1001270402.

Riedl, J. et al. (2008) ‘Lifeact: A versatile marker to visualize F-actin’, Nature Methods, 5(7), pp. 605–607. doi: 10.1038/NMETH.1220.

Rocchetti, A. et al. (2014) ‘Fluorescent labelling of the actin cytoskeleton in plants using a cameloid antibody’, Plant Methods, 10(1), p. 12. doi: 10.1186/1746-4811-10-12.

Rochefort, N. L. and Konnerth, A. (2012) ‘Dendritic spines: from structure to in vivo function.’, EMBO reports. Nature Publishing Group, 13(8), pp. 699–708. doi: 10.1038/embor.2012.102.

Rudy, J. W. (2015) ‘Actin dynamics and the evolution of the memory trace’, Brain Research. Elsevier, 1621, pp. 17–28. doi: 10.1016/j.brainres.2014.12.007.

Schubert, V. and Dotti, C. G. (2007) ‘Transmitting on actin: synaptic control of dendritic architecture.’, Journal of cell science, 120(Pt 2), pp. 205–212. doi: 10.1242/jcs.03337.

Star, E. N., Kwiatkowski, D. J. and Murthy, V. N. (2002) ‘Rapid turnover of actin in dendritic spines and its regulation by activity’, Nature Neuroscience, 5(3), pp. 239–246. doi: 10.1038/nn811.

Tárnok, K. et al. (2015) ‘A new tool for the quantitative analysis of dendritic filopodial motility’, Cytometry Part A, 87(1), pp. 89–96. doi: 10.1002/cyto.a.22569.

Traenkle, B. and Rothbauer, U. (2017) ‘Under the microscope: Single-domain antibodies for live-cell imaging and super-resolution microscopy’, Frontiers in Immunology. Frontiers Media S.A., 8(AUG), p. 1030. doi: 10.3389/FIMMU.2017.01030/BIBTEX.

Wegner, W. et al. (2017) ‘In vivo mouse and live cell STED microscopy of neuronal actin plasticity using far-red emitting fluorescent proteins’, Scientific reports. Sci Rep, 7(1). doi: 10.1038/S41598-017-11827-4.

Xu, R. and Du, S. (2021) ‘Overexpression of Lifeact-GFP Disrupts F-Actin Organization in Cardiomyocytes and Impairs Cardiac Function’, Frontiers in Cell and Developmental Biology. Frontiers Media S.A., 9. doi: 10.3389/FCELL.2021.746818/FULL.

Yuste, R. (2010) Dendritic spines, MIT Press. doi: 10.7551/mitpress/9780262013505.001.0001.

Yuste, R. and Bonhoeffer, T. (2004) ‘Genesis of dendritic spines: insights from ultrastructural and imaging studies.’, Nature reviews. Neuroscience, 5(1), pp. 24–34. doi: 10.1038/nrn1300.

